# Sex differences in diverse conditioned fear behaviors following systemic naloxone administration

**DOI:** 10.64898/2026.08.20.745978

**Authors:** EM Greiner, ML Laine, J Fourte, RM Shansky

## Abstract

Fear conditioning studies have historically relied on freezing as the primary measure of conditioned fear despite evidence that defensive responding is behaviorally diverse and sexually dimorphic. The endogenous opioid system, particularly mu-opioid receptor (MOR) signaling, is known to regulate fear learning and conditioned analgesia, yet its role in alternative fear-related behaviors and sex differences remains unclear. Here, we investigated the effects of systemic naloxone administration prior to auditory fear conditioning on freezing, darting, shock responsivity, and ultrasonic vocalizations (USVs) in male and female rats. Adult Sprague Dawley rats received naloxone (5 mg/kg, i.p.) or saline prior to conditioning and underwent fear recall testing 24 hours later. Naloxone produced sex- and behavior-specific effects across conditioning and recall. During conditioning, naloxone increased freezing in males during baseline and early tone presentations, while females exhibited reduced shock-response velocity and increased post-shock freezing. Naloxone did not significantly alter darting or USV production during conditioning. During recall, freezing behavior did not differ across groups. Naloxone-treated females, however, exhibited a distinct alarm-calling pattern, with fewer callers overall but increased call output among those that vocalized. These findings suggest that MOR antagonism differentially alters distinct components of fear expression in a sex-dependent manner and support the idea that freezing and alarm calling may reflect separable aspects of fear processing.

## [1] Introduction

Pavlovian fear conditioning is a powerful tool to investigate the neural circuitry involved in fear learning and threat responses. In this paradigm animals learn to associate a previously neutral cue, such as an auditory tone (conditioned stimulus; CS) with an aversive event, typically a foot shock (unconditioned stimulus; US). Repeated paired presentations of the CS followed by US eventually result in the animal displaying a conditioned fear response (CR) to the CS alone. Experimenters are then able to determine the degree of fear learning and strength of the subsequent fear memory through measuring the CR. Freezing is the most common CR measured during fear conditioning, defined as the lack of all movement except that required by respiration (Fanselow, 1980; Fanselow, 1994).

Two main issues arise in the classical applications of this paradigm in pre-clinical models. First, using freezing as the sole measure of learning does not capture the full diversity of behavioral responses to fear (see Chu et al., 2024). Second, much of this work has focused solely on males, often excluding females entirely (Beery & Zucker, 2011; Zucker et al., 2022). These factors limit our understanding of the full range of individual variation in fear behavior and the underlying neurobiological mechanisms that mediate fear learning and memory.

Our lab has identified an alternative, active, conditioned fear response that is exhibited predominantly by females, known as darting (Gruene et al., 2015; Mitchell et al., 2022). Darting is characterized by rapid movements across the conditioning chamber during the presentation of the CS. Another commonly expressed threat response during fear conditioning is the emittance of ultrasonic vocalizations (USVs), particularly those within the 22hz frequency deemed ‘alarm calls’ (Wöhr et al., 2005; Schwarting et al., 2007; Dupin et al., 2019). Much work has found that males produce more alarm calls (22 kHz) than females throughout conditioning and testing (Graham et al., 2009; Laine et al., 2022; Schwarting, 2018).

Interestingly, both behaviors are related to sex differences in unconditional responses. Female darters exhibit an increased shock response velocity (how quickly the animal moves in response to the shock) (Gruene et al., 2015; Mitchell et al., 2022), and increased shock intensity is positively correlated to the number of emitted alarm calls in males only (Kosten et al., 2006; Laine et al., 2022).

The demonstrated variability in shock response may be related to sex differences in pain perception. Fear conditioning is known to induce fear conditioned analgesia (Fanselow, 1986; Fanselow & Helmstetter, 1988; Helmstetter, 1993), in which exposure to the CS results in increased endogenous opioid activity (Chance et al, 1978) to attenuate the aversive impact of the resulting US. However, several studies have found that females show little to no fear conditioned analgesia following conditioning (Llorente-Berzal et al., 2022; Stock et al., 2001). Additionally, females are more sensitive to noxious stimuli than males, who show a higher responsiveness to mu-opioid agonists (Wiesenfeld-Hallin, 2005).

Naloxone is a commonly used mu-opioid receptor (MOR) antagonist. Although non-specific, it has the highest affinity for MORs (Martin, 1967; Martin, 1976). Previous work has found that administration of the naloxone facilitates fear acquisition (measured by freezing) to both discrete cues and context (Fanselow & Bolles, 1979a). This effect has been observed in both males and females separately (Fanselow & Bolles, 1979b; McNally et al., 2004). However, investigation of differences between the sexes, or of the effect of mu-opioid antagonism on alternative behavioral expressions of fear has not been done. The present study aims to determine the impact of systemic naloxone administration prior to conditioning on both fear learning and memory as measured by freezing, darting, and the emittance of ultrasonic vocalizations for both males and females.

## [2] Materials & Methods

### [2.1] Subjects

Adult male (24) and female (24) Sprague Dawley rats (Charles River Laboratories) that weighed 250-300g for females and 350-400g for males were pair housed and maintained on a 12-hour light/dark cycle (lights on: 7:00 AM) in a temperature (22°C ±1°C) and humidity (40% ±10%) controlled colony room. Males and females were housed in the same colony room and given *ad libitum* access to water and standard rat chow (RMH 3000, Purina). Animals were given one week to acclimate to the colony housing room prior to behavioral testing. Each cage contained a tinted Plexiglas chamber for nesting and enrichment, and heat-treated pine shavings for bedding. All behavioral experiments were conducted during the light phase between 9:00 A.M. and 4:00PM. All housing and testing procedures were in compliance with the National Institutes of Health Guidelines for Care and Use of Laboratory Animals and approved by the Northeastern University Institutional Animal Care and Use Committee.

Rats were divided evenly (n=12) into four variable groups: female vehicle, female naloxone, male vehicle, and male naloxone. Three males and one female were excluded from final analysis due to issues with data collection. The resulting final n’s were as follows: female vehicle n=11, female naloxone n=12, male vehicle n=10, male naloxone n=11.

### [2.2] Behavioral data collection

#### [2.2.1] Experiment set up

All animals were handled during the 2 days prior to fear conditioning for 5 minutes each. They were habituated to transport from the vivarium to the testing room on a cart the day prior to conditioning. On the conditioning day, animals were brought into the testing room 30 minutes before the start to habituate to the testing room and ambient noise. Following habituation, they were given an i.p. injection of either naloxone or saline and left for fifteen minutes to allow for full efficacy. They were then placed in sound-attenuating conditioning chambers (model H1024A Rat Test Cage, Coulbourn Instruments), consisting of Plexiglas, metal walls, and a metal grid floor for the delivery of footshocks (model H10-11R-TC Shock Floor, Coulbourn Instruments).

The chambers were dimly lit by an overhead light (2lux). Following a 5 min baseline period with no stimulus presentations, the rats were sequentially exposed to a total of seven CS–US pairings (Fig. 1A). The CS used a 30 second 4kHz tone played in each chamber by a speaker measuring between 73-80dB as measured at the center of the chamber. During conditioning the tone co-terminated with a 0.5 second 0.5mA footshock. The intertrial interval (ITI) was of varying lengths between each of the CS–US pairings (90–330s). Two to four rats were conditioned simultaneously in the same room, without mixing sexes within each run. Chambers and the testing cages were cleaned with water and ethanol between each run.

**Figure 1.**
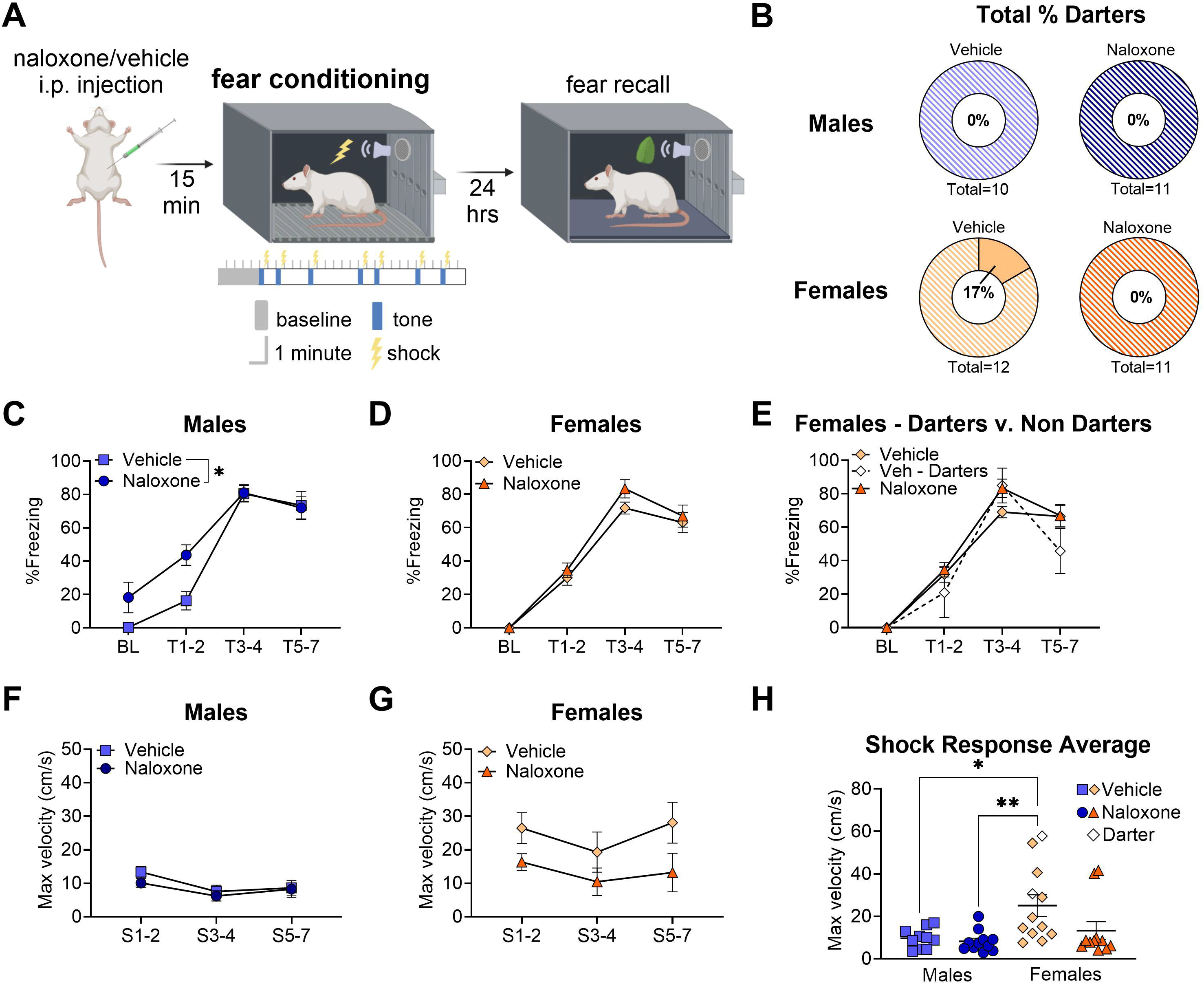
Naloxone alters unconditioned behaviors in males and females during fear conditioning. **A**. Graphical representation of the timing and parameters of fear conditioning. **B**. Pie charts showing the proportion of darters (solid color) and non-darters (striped color) across each experimental group (top: males, bottom: females). **C-D**. Line graph showing the percentage of time spent freezing during each tone bin for animals injected with naloxone and injected with a vehicle prior to conditioning (BL: baseline, T1-2: tones 1 and 2, T3-4: tones 3 & 4, T5-7: tones 5, 6, & 7). **E**. Line graph showing the percent of time spent freezing during each tone bin by female groups with the vehicle injected group split into darters and non-darters. **F-G**. Line graph showing the shock response velocity (how quickly the animal moved in response to the footshock) for each shock bin for animals injected with naloxone and injected with a vehicle prior to conditioning (S1-2: shocks 1 & 2, S3-4: shocks 3 & 4, S5-7: shocks 5, 6, & 7). **H**. Bar graph showing the average shock response velocity across conditioning for each experimental group. All graphs depict the mean ± SEM and each dot on the bar graph represents a single animal. Significant main effects and post hoc comparisons are denoted with asterisks depicting degree of significance (1: p<0.05, 2: p<0.01)

Fear recall testing occurred 24 hours after conditioning. Animals were brought into the testing room 30 minutes before the start of the test to habituate. The testing room and chambers were the same as for fear conditioning, but with different lighting (8lux), scent (2–4 drops of Dr Bronner’s peppermint-scented pure-castile liquid soap placed on a train underneath the test cage floor) and chamber features (black Plexiglas floor covering the metal grids). Animals were exposed to a baseline period of 5 minutes followed by three presentations of the same tone, in the absence of footshock, at varying intervals (150–240 s) (Fig. 3A).

We recorded rodent calls ranging from 0-120kHz during conditioning and analyzed the calls within three distinct call categories: baseline, shock, & alarm. During the 5-minute baseline period at the start of conditioning (Fig. 2A), rodents emitted various calls typically ranging from 40-60kHz (Fig. 2B), from here on referred to as ‘baseline calls’. Previous work in our lab has found that females, on average, emit a greater number of these than males (Laine et al., 2022). Despite vehicle females having a slightly elevated number of these calls, we found only a trending effect of experimental group in baseline calls emitted during our fear conditioning session (Fig. 2D) (*One-way ANOVA: trending group effect, F(3,40)=2.472, p=0.076*).

**Figure 2.**
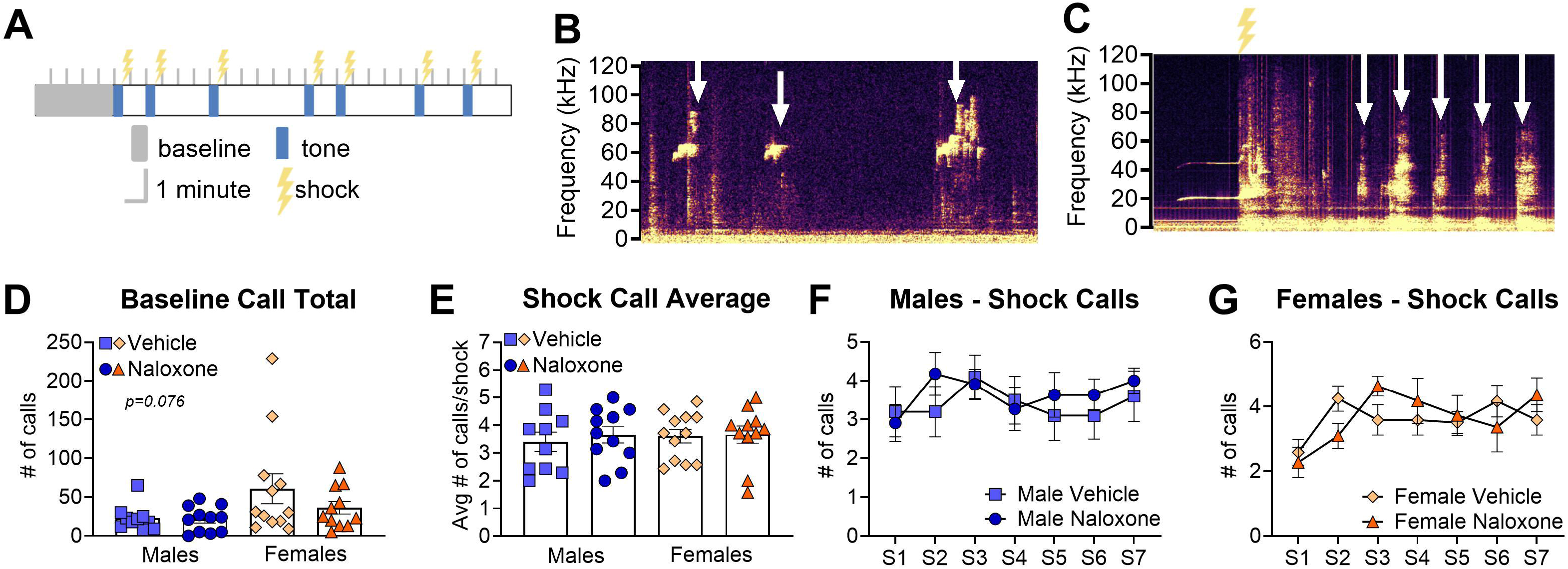
Naloxone does not affect the emittance of baseline or shock calls during fear conditioning. **A**. Graphical representation of the timing of event related epochs during fear conditioning. **B**. Representative spectrogram (from DeepSqueak) showing typical high- frequency ultrasonic calls (white arrows) recorded during the baseline period. **C**. Representative spectrogram showing typical shock calls (white arrows). Yellow lightning symbol denotes shock onset time. **D**. Bar graph depicting the total baseline calls emitted during the first five minutes of the conditioning session for each experimental group. **E**. Bar graph depicting the average number of shock calls emitted per shock for each experimental group. **F-G**. Line graphs depicting the shock calls emitted for each shock (S) for males and females injected with naloxone and injected with a vehicle prior to conditioning. All graphs depict the mean ± SEM and each symbol on the bar graphs represents a single animal. Trending main effect denoted with p value.

**Figure 3.**
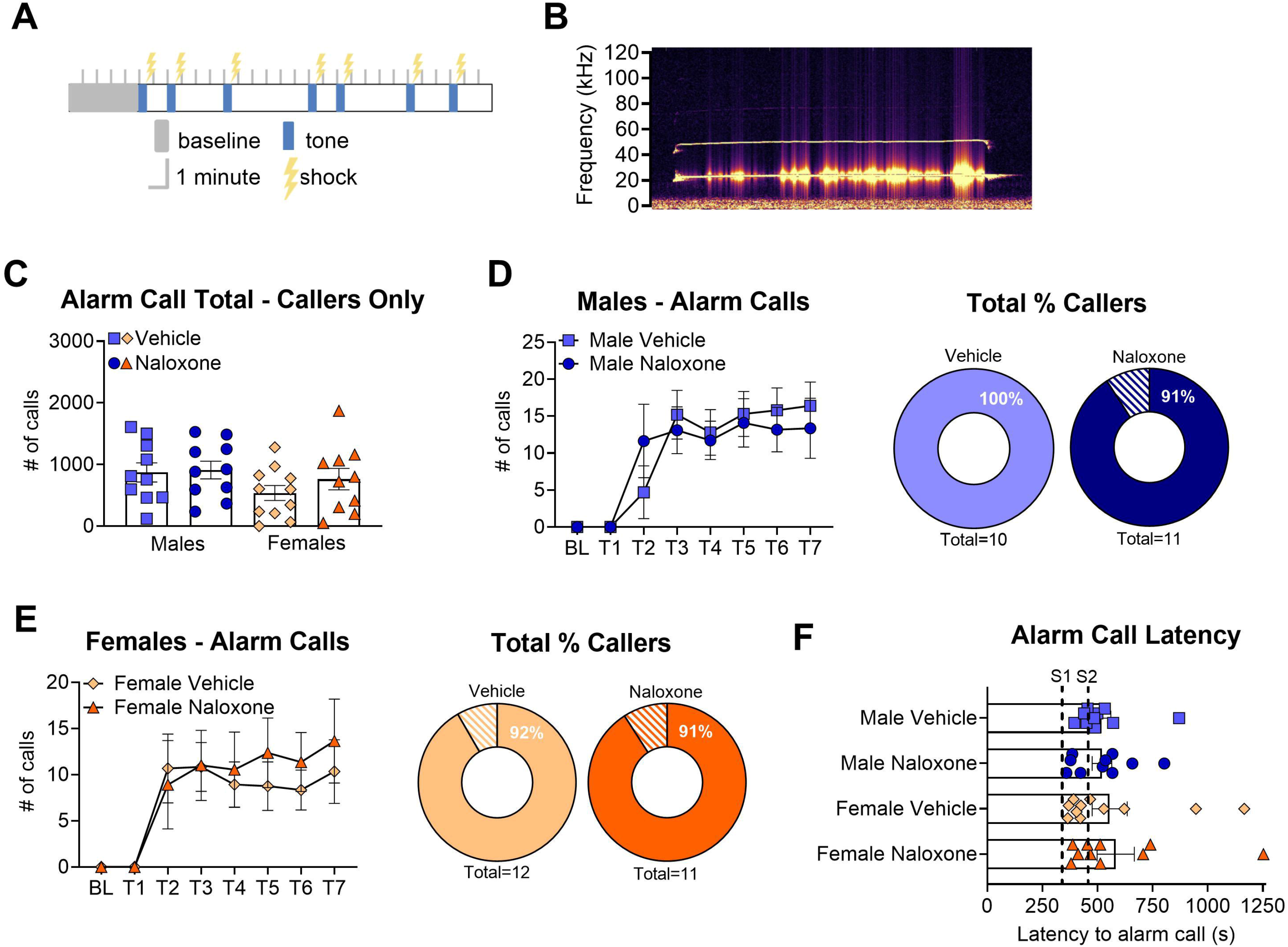
Naloxone does not affect the emittance of 22kz alarm calls during fear conditioning **A.** Graphical representation of the timing of event related epochs during fear conditioning. **B.** Representative spectrogram (from DeepSqueak) showing an ultrasonic alarm call captured during an intertrial interval, with a principal frequency of ∼22kHz. **C**. Bar graph depicting total number of emitted alarm calls for each experimental group. Graph only depicts animals that emitted at least one alarm call during the conditioning session. **D**. Left: Line graph depicting the number of calls emitted during each tone by males that were injected with naloxone or vehicle prior to conditioning (BL: baseline, T: tone). Right: Pie charts showing the proportion of callers (solid color) and non-callers (striped color) during conditioning for naloxone and vehicle treated males. **E**. Left: Line graph depicting the number of calls emitted during each tone by females that were injected with naloxone or vehicle prior to conditioning. Right: Pie charts showing the proportion of callers (solid color) and non-callers (striped color) during conditioning for naloxone and vehicle treated females. **F**. Bar graph depicting the latency to emit the first alarm call for animals in each experimental group. The first dotted line denotes the presentation of the first footshock (S1) and the second dotted line denotes the presentation of the second footshock (S2). All graphs depict the mean ± SEM and each symbol on the bar graphs represents a single animal

#### [2.2.2] Naloxone administration

Animals were weighed the morning of testing to determine their body weight. Fifteen minutes prior to fear acquisition males and females in naloxone groups received an i.p. injection at a dosage of 5mg/kg of 5 mg/ml Naloxone HCL (Sigma-Aldritch, St. Louis, MO) dissolved in 0.9% sterile saline. A dosage of 5mg/kg represents the moderate to high dosage range previously reported for naloxone i.p. injections (Fanselow & Bolles, 1979a; Fanselow & Bolles, 1979b). Males and females in the vehicle group received an i.p. injection of 0.9% sterile saline, also proportional to their body weight.

#### [2.2.3] Behavior tracking

Stimulus delivery was controlled, and videos of the animals’ behavior were recorded using Ethovision (version 16; Noldus Technologies) and infrared digital cameras mounted on top of each conditioning chamber. Time spent freezing, the occurrence of darts, and the maximum velocity the rat reached as a response to the shock were analyzed using freely available Python-based software ScaredyRat (Mitchell et al., 2022). An animal was classified as a Darter if it performed one or more darts (movement at a speed exceeding 20cm/s) during the presentation of the tone (excluding the first two tones)(Mitchell et al., 2022; Mitchell et al., 2024). Baseline freezing was recorded from the first 2 minutes of the 5-minute stimulus-free time in the chamber at the start of the session. Freezing was defined in ScaredyRat as the absence of observable movement, with a minimum bout duration of 1 second.

#### [2.2.4] Ultrasonic vocalization recording

Vocalizations emitted in the audible and ultrasonic range (0–120kHz; sampling rate, 250kHz) were recorded using microphones (model CM16, Avisoft Bioacoustics) mounted over each conditioning chamber. The audio files were processed with DeepSqueak (version 3; Coffey et al., 2019), a publicly available user interface that uses machine learning to detect spectrograms typical of rodent USVs. All detected calls were then manually confirmed by a trained investigator. Temporal alignment of the audio and behavioral data was confirmed by observation of the tone cue within the audible range of the spectrogram. We then aligned each detected call with an epoch (baseline, tone, shock, or ITI) and identified alarm calls based on specific criteria (call length, 70ms; main frequency of the call, 30kHz; change in frequency, 10kHz).

### [2.3] Statistical analysis

Group differences were analyzed using Graphpad Prism (version 11). Analysis of behavioral data (freezing, shock response, ultrasonic vocalizations) across tone or shock presentations was analyzed using a 2-way repeated measures ANOVA to compare the effect of the drug across both conditioning and testing within male and female groups. Geisser– Greenhouse correction for nonsphericity was applied when needed. Difference in the number of darters within each group and the number of animals who emitted 22kHz ‘alarm calls’ during both fear conditioning and recall was determined using a Fisher’s Exact Test. Analysis of average behavioral responding (shock response & ultrasonic vocalizations) between the sexes was conducted using a one-way ANOVA for both fear conditioning and fear recall. All *post hoc* multiple comparisons were Bonferroni corrected.

## [3] Results

### [3.1] Fear acquisition

A graphical representation of the timing and parameters of fear conditioning can be found in Fig. 1A.

#### [3.1.1] Naloxone alters unconditioned behaviors in males and females during fear conditioning

Each animal was classified as a ‘Darter’ or ‘Non-darter’ based on whether they performed at least one dart (movement exceeding velocity of 20 cm/s) during the presentation of tones 3-7 (Gruene et al., 2015; Colom-Lapetina et al., 2019; Mitchell et al., 2022). We observed no significant difference in the number of Darters between the sexes or drug groups (*Fisher’s Exact, p=0.23*). In fact, only two rats, both in the female vehicle group, exhibited darting behavior during conditioning. This is a low number of Darters compared to typical darting rates observed in our lab, which tend around 40% for females and 10% for males (Gruene et al., 2015).

Analysis of freezing behavior throughout conditioning revealed that males administered naloxone exhibited higher rates of freezing compared to vehicle males, largely concentrated to the baseline period and early conditioning (tones 1 & 2) (Fig. 1C) (*2-way RM ANOVA: main effect of drug, F(1,19)=4.49, p=0.048; main effect of tone, F(3, 57)=65.18, p<0.0001; trending tone x drug interaction, F(1.914, 36.37)=2.712, p=0.082*). Females, on the other hand, did not show any drug effect on freezing behavior, with both naloxone administered females and vehicle females exhibiting similar freezing levels across conditioning (Fig. 1D) (*2-way RM ANOVA: main effect of tone, F(3,63)=139.8, p<0.0001; no main effect of drug, F(1,21)=1.789, p=0.2; no tone x drug interaction, F(3,63)=0.6663, p=0.58*). Female Darters, additionally, did not freeze significantly differently compared than Non-darters in either group (Fig. 1E) (*2-way RM ANOVA: main effect of tone, F(2.337,46.74)=83.37P, p<0.0001; no main effect of group, F(2,20)=1.021, p=0.38; no tone x group interaction, F(4.674,46.74)=1.438, p=0.23*). Previously our lab found that Darters freeze less than Non-darters (Gruene et al., 2015), however, the null findings here are likely the result of the low number of Darters within this experiment.

Shock response velocity (speed at which the rat moves in direct response to receiving a footshock) followed more typical patterns for what we have previously observed within the control groups. The female vehicle group had a greater average shock response velocity compared to both male vehicle and male naloxone groups (Fig. 1H) (*one-way ANOVA: main effect of experimental group, F(3,40)= 2.809, p= 0.007; Bonferroni post hoc comparison: female vehicle v. male vehicle p=0.01, female vehicle v. male naloxone p= 0.027*). This same difference was not observed for the female naloxone group (*Bonferroni post hoc comparison: female naloxone vs. male vehicle & male naloxone p>0.99*). We have previously observed a marked sex-dependent difference in shock response, with females exhibiting velocities approximately 15 cm/s higher than their male counterparts (Mitchell et al., 2022), making the female naloxone shock response rates closer to that typically observed in males. However, no significant drug effects were observed within male or female groups across shock presentations (Fig. 1F-G) (*2- way RM ANOVA, no main effect of drug in males: F (1,19)=0.7037, p=0.4; no main effect of drug in females: F(1,21)=3.121, p=0.09*)

The difference in shock response in females given naloxone and in early freezing behavior for males given naloxone, suggest that MOR receptor activity may have a greater impact on unconditioned behaviors during fear acquisition.

#### [3.1.2] No effect of Naloxone on ultrasonic vocalizations during fear conditioning

We additionally quantified a specific call that is observed during the presentation of the footshock (Fig. 2C), which has been previously identified and defined in Laine et al., 2022, referred to as ‘shock calls’. There was no significant difference between drug groups in the number of shock calls emitted across conditioning for males or females (Fig. 2E-F) (*2-way RM ANOVA: no main effect of drug in males F(1,19)=0.3013, p=0.6; no effect of drug in femalesF(1,21)=0.01971, p=0.9*). Additionally, males did not have a significant difference in the number of calls emitted per shock across the conditioning session (*no effect of shock number, F(6,114)=0.8863, p=0.5*). Females, on the other hand, emitted a greater number of shock calls during the 2^nd^, 3^rd^, and 7^th^ presentation of the shock compared to the first (*main effect of shock number, F(3.211,67.42)=3.155, p=0.028; Bonferroni post hoc comparison: S1 v. S2 p= 0.006, S1 v. S3 p= 0.01, S1 v. S7 p= 0.008*). However, there was also no overall difference in the average number of calls per shock between groups (*One-way ANOVA: no main effect of group, F(3,40)=0.1564, p=0.9*), with most animals emitting 3-4 calls per shock (Fig. 2G).

The third call type that we quantified were 22kHz ‘alarm calls’ (Fig. 3B) that have been previously established as a distinct type of ultrasonic vocalization emitted in response to aversive events and threats (Burgdorf et al., 2000; Wöhr et al., 2005; Borta et al., 2006; Yee et al., 2012; Fendt et al., 2018) and are commonly emitted during fear conditioning. In each group almost all animals (90-100%) emitted at least one alarm call during conditioning (Fig. 3D&E), with no significant difference in the number of ‘callers’ within each group (*Fisher’s Exact, p>0.99*). All animals increased alarm calling across conditioning (*2-way RM ANOVA, main effect of tone in males F(7,133)=17.55, p<0.0001; main effect of tone in females F(3.569,74.94)=9.769, p<0.0001*). However, we did not observe any differences between drug groups in the number of calls emitted across tone presentations (*no main effect of drug in males F(1,19)=0.01601, p=0.9; no main effect of drug in females F(1,21)=0.1637, p=0.7*) or in the total number of alarm calls emitted during the entire conditioning session (Fig. 3C) (*one-way ANOVA: no main effect of experimental group, F(3,40)=1.324, p=0.3*). There was also no significant difference between groups in the latency to emit their first alarm call (*one-way ANOVA: no main effect of experimental group, F(3,32)=0.8949, p=0.5*), with most animals starting to alarm call following the second footshock (Fig. 3F). Overall, there was no identifiable effect of blocking MORs on vocalizations during conditioning.

#### [3.1.2] Naloxone increases post-shock freezing in females

To examine the effect of naloxone administration on behavior during non-cue periods we examined freezing during the post-shock period (PS) (30 seconds following the footshock) and alarm calling across each ITI period.

Males did not show any effect of naloxone on the amount of time spent freezing post-shock (Fig. 4B) (*2-way RM ANOVA: no main effect of drug, F(1,19)=0.2748, p=0.61; no drug x post-shock period interaction, F(4.387,83.34)=0.6464, p=0.65*) or in the number of alarm calls emitted during the ITI periods (Fig. 4E) (*2-way RM ANOVA: no main effect of drug, F(1,19)=0.04487, p=0.83; no drug x ITI period interaction, F(2.214,42.06)=0.4052, p=0.69*). There was a general increase in both behaviors across the conditioning session (*freezing: main effect of post-shock period, F(4.387,83.34)=15.76, p<0.0001; alarm calls: main effect of ITI period, F(2.214,42.06)=37.14, p<0.0001*).

**Figure 4.**
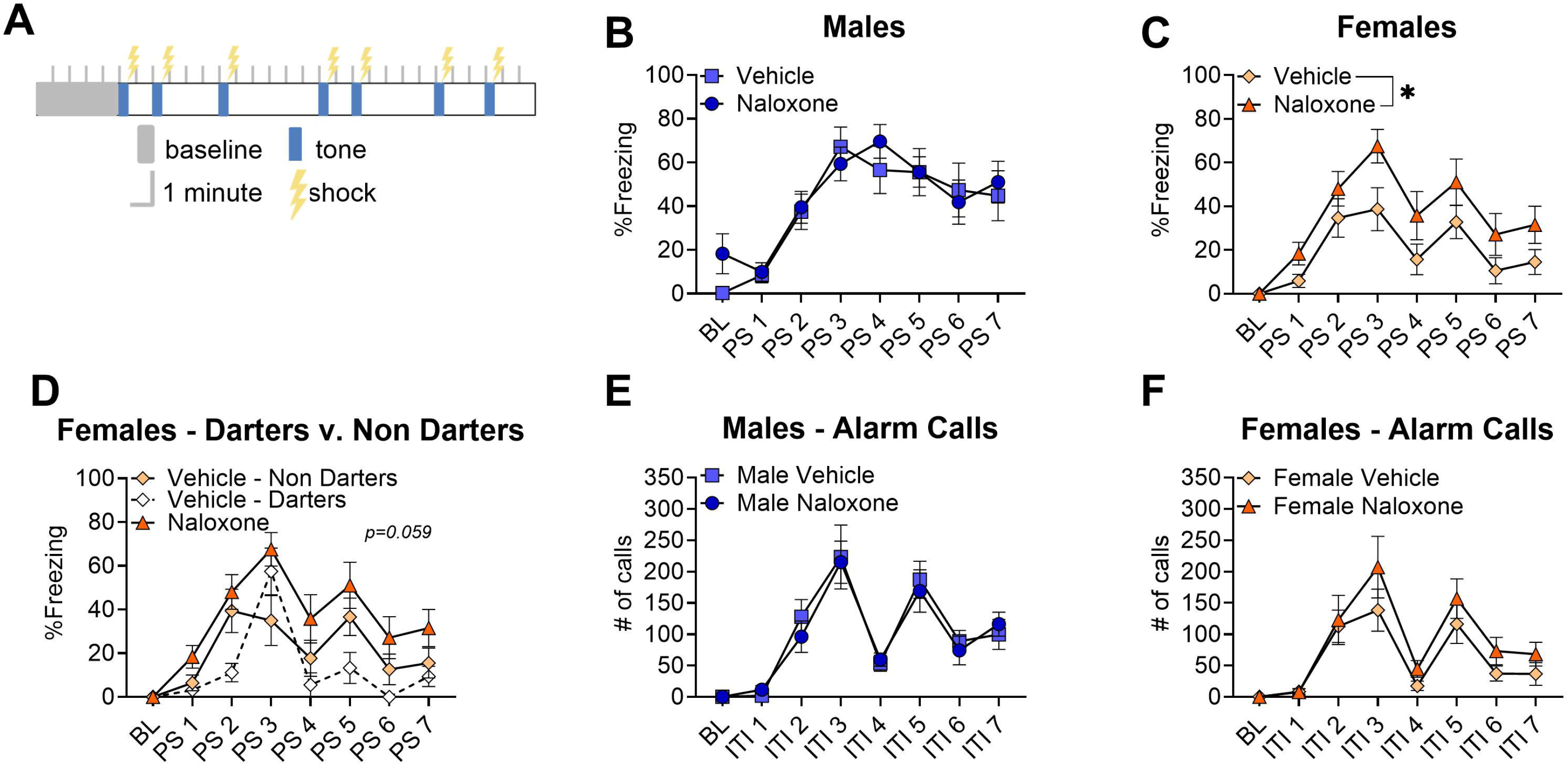
Naloxone increases post-shock freezing in females **A**. Graphical representation of the timing of event related epochs during fear conditioning. **B-C**. Line graphs showing the percentage of time spent freezing during each post-shock period (30s following footshock) for male and female animals injected with naloxone and injected with a vehicle prior to conditioning (BL: baseline, PS: post-shock). **D**. Line graph showing the percent of time spent freezing during each post-shock period by female groups with the vehicle injected group split into darters and non-darters. **E-F**. Line graphs showing the number of alarm calls emitted during each ITI period for male and female animals injected with naloxone and injected with a vehicle prior to conditioning (BL: baseline, ITI: intertrial interval). All graphs depict the mean ± SEM. Significant main effects are denoted with asterisks depicting degree of significance (1: p<0.05)

Females, additionally did not show any differences in number of emitted alarms calls during ITIs, but showed a general increase in calling across the session (Fig. 4F) (*2-way RM ANOVA: main effect of ITI period, F(2.320,44.08)=22.74, p<0.0001; no main effect of drug, F(1,19)=1.376, p=0.26; no drug x ITI period interaction, F(2.320,44.08)=0.7680, p=0.49*).

However, females given naloxone froze at higher rates than vehicle females during the post- shock period (Fig. 4C) (*2-way RM ANOVA: main effect of drug, F(1,21)=6.263, p=0.021; main effect of post-shock period, F(4.551,95.57)=15.11, p<0.0001; no drug x post-shock period interaction, F(4.551,95.57)=0.8167, p=0.53*). When Darters were analyzed as a separate group we no longer see a significant effect of group, however there is a trending group difference (Fig. 4D) (*F(2, 20)=3.269, p=0.059*), suggesting that the lower freezing rates in the vehicle group are being enhanced by the presence of Darters. This replicates previous findings that naloxone increases post-shock freezing rates in females specifically (Fanselow & Bolles 1979a).

### [3.2] Fear recall

A graphical representation of the timing and parameters of fear recall can be found in Fig. 5A.

**Figure 5.**
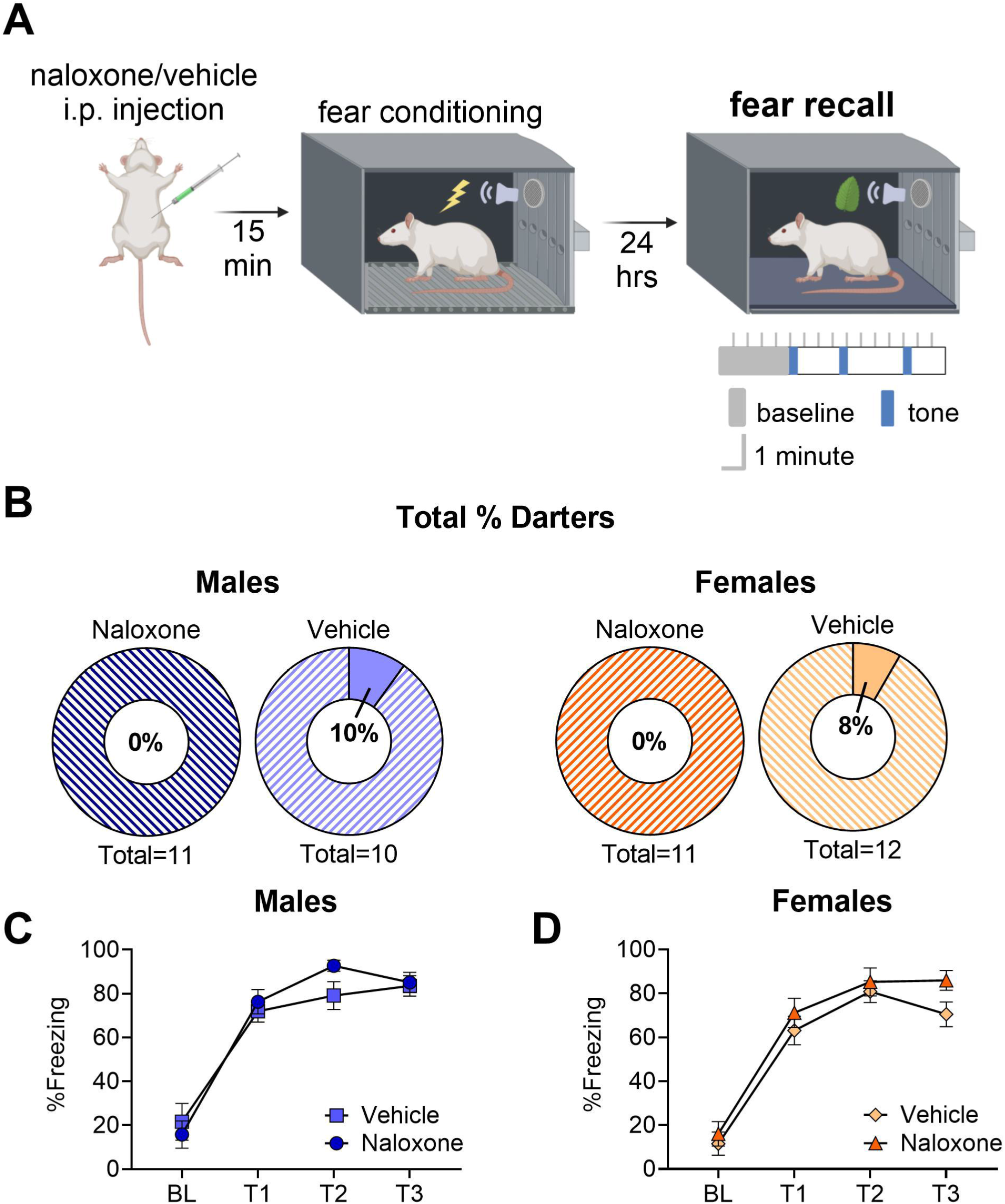
No effect of naloxone on freezing or darting during fear recall **A**. Graphical representation of the timing and parameters of fear recall. **B**. Pie charts showing the proportion of darters (solid color) and non-darters (striped color) across each experimental group (left: males, right: females). **C-D**. Line graphs showing the percentage of time spent freezing during each tone for animals injected with naloxone and injected with a vehicle prior to conditioning (BL: baseline: T: tone). All graphs depict the mean ± SEM

#### [3.2.1] No effect of naloxone on freezing or darting during fear recall

Darting was observed in one animal in each of the vehicle groups and not observed in either of the naloxone groups (Fig. 5B). A Fisher’s Exact test yielded a non-significant result, confirming no difference in the number of Darters per group (*p= 0.6*). This aligns with previous work in our lab that has found substantially decreased rates of darting during recall compared to conditioning (Mitchell et al., 2022).

Freezing rates indicate that all animals successfully remembered the tone-shock association (Fig. 5C&D) (*2-way RM ANOVA: main effect of tone in males, F(1.992,37.84)=75.97, p<0.0001, Bonferroni post hoc comparison of baseline vs. T1, T2, & T3 all p<0.001; main effect of tone in females F(2.324,48.80)=84.31, p<0.0001, Bonferroni post hoc comparison of baseline vs. T1, T2, & T3 all p<0.001*), and time spent freezing during each tone did not differ based on drug group for either males or females (*no main effect of drug in males F(1,19)=0.5327, p=0.5; no main effect of drug in females F(1,21)=2.220, p=0.1511*). Overall, blocking of MORs prior to fear conditioning did not appear to alter freezing or darting behavior during the tone.

#### [3.2.2] Naloxone alters alarm calling in females during fear recall

Female rats in the vehicle group emitted slightly more baseline calls than any other group (Fig. 6B). A one-way ANOVA revealed a trending effect of experimental group on the number of emitted baseline calls (*F(3,40)=2.444, p=0.074*). Because the fear recall test omits the footshock, the only other call quantified was alarm calls. There was a noticeable decrease in the number of naloxone females that emitted alarm calls during the testing session, with only 55% emitting at least one call (Fig. 6E). A Fisher’s Exact test revealed that this was a significantly different proportion of callers and non-callers compared to all other testing groups (Fig. 6D) (*Fisher’s Exact, p=0.04*), whose rates of callers are comparable to fear conditioning.

**Figure 6.**
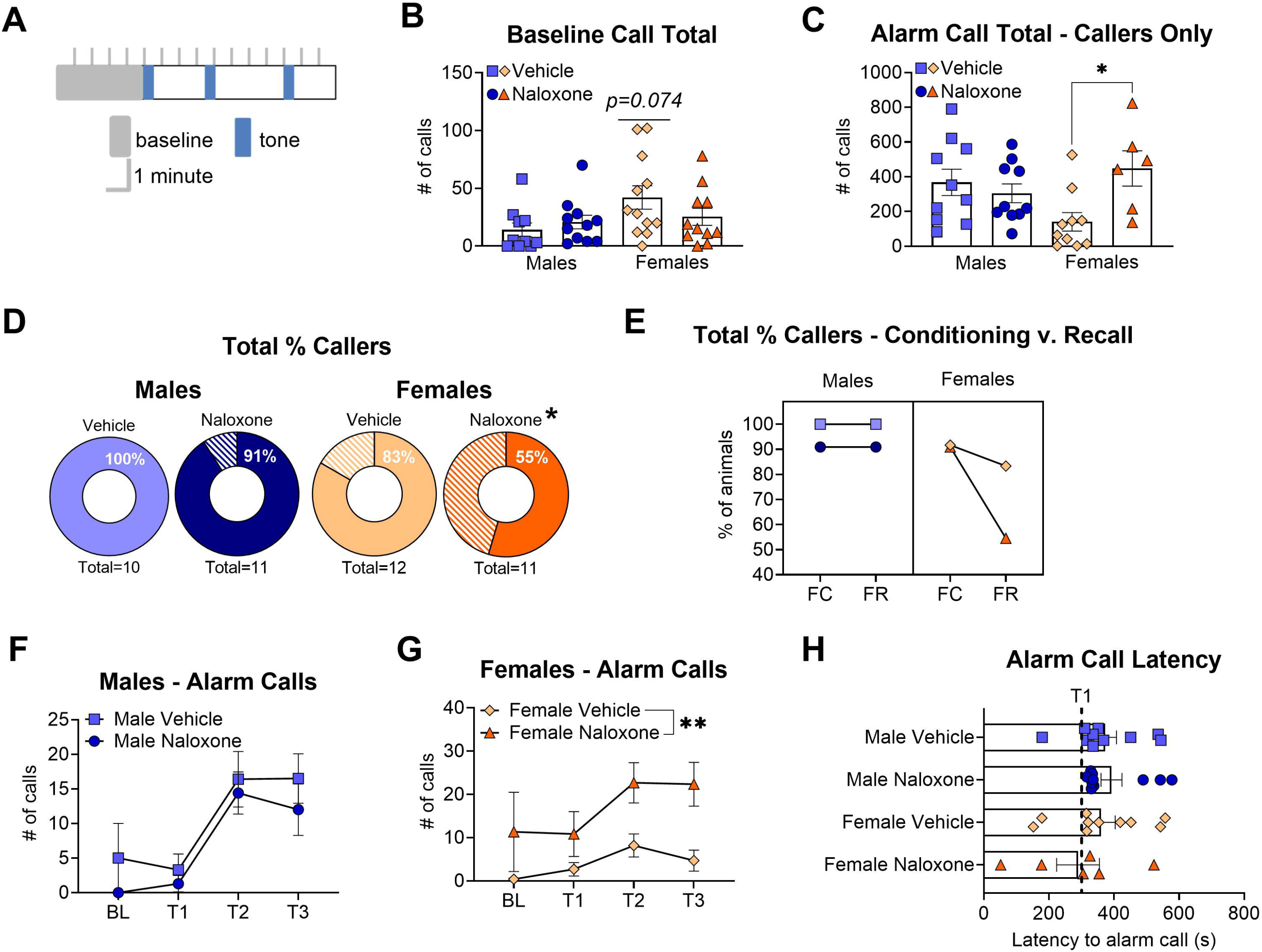
Naloxone alters alarm calling in females during fear recall. **A**. Graphical representation of the timing of event related epochs during fear recall. **B**. Bar graph depicting the total baseline calls emitted during the first five minutes of the conditioning session for each experimental group. **C**. Bar graph depicting total number of emitted alarm calls for each experimental group. Graph only depicts animals that emitted at least one alarm call during the testing session. **D**. Pie charts showing the proportion of callers (solid color) and non-callers (striped color) during fear recall for naloxone and vehicle treated males (left) and females (right). **E**. Line graphs depicting the change in prorportion of callers from fear conditioning to fear recall (males left, females right). **F-G**. Line graphs depicting the number of calls emitted during each tone by males and females that were injected with naloxone or vehicle prior to conditioning. **H**. Bar graph depicting the latency to emit the first alarm call for animals in each experimental group. The first dotted line denotes the presentation of the first tone (T1). All graphs depict the mean ± SEM and each dot on the bar graph represents a single animal. Significant main effects and post hoc comparisons are denoted with asterisks depicting degree of significance (1: p<0.05, 2: p<0.01). Trending main effect denoted with p value

Despite this difference, there was no significant drug effect on the number of alarm calls emitted across the tone presentations for either females or males (Fig. 6F&G) (*2-way RM ANOVA: no main effect of drug in males F(1,19)=1.598, p=0.2; no main effect of drug in females F(1,21)=2.741, p=0.1*). There was a significantly greater number of total emitted alarm calls during the testing session for females given naloxone compared to vehicle females (Fig. 6C) (*one-way ANOVA: main effect of experimental group, F(3,32)=3.419, p=0.029; Bonferroni post hoc comparison naloxone female v. vehicle female p= 0.04*). Additionally, naloxone females had a shorter latency to emit their first alarm call (Fig. 6H), but there was no significant difference in the overall latency to call between groups (*one-way ANOVA: no main effect of experimental group, F(3,32)=0.8949, p=0.5*). Taken together, these results suggest that blocking MORs prior to fear conditioning alters the pattern of alarm call emission during fear recall in females.

#### [3.2.3] Naloxone increases ITI alarm calling during fear recall in females

To examine the effect of naloxone administration on behavior during non-cue periods we examined freezing during the pre-tone period (PT) (30 seconds prior to tone onset) and alarm calling across each ITI period.

Males and females given naloxone did not differ from vehicle groups in their rates of freezing during the pre-tone period (Fig. 7B&C) (*2-way RM ANOVA: no main effect of drug in males, F(1,19)=0.9024, p=0.35; no main effect of drug in females, F(1,21)=1.886, p=0.18*). We additionally did not observe any differences between female Darters and Non-darters (Fig. 7D) (*2-way RM ANOVA: no main effect of group, F(2,20)=0.9137, p=0.42*). Notably, all animals exhibited unexpectedly high rates of freezing following the first tone presentation (PT 2 & 3) (*2-way RM ANOVA: main effect of pre-tone period in males, F(2.578,48.99)=77.9, p<0.0001, Bonferroni post hoc comparison BL v. PT2 & PT3 all p<0.0001; main effect of pre-tone period in females, F(1.567, 32.92)=84.26, p<0.0001, Bonferroni post hoc comparison BL v. PT2 & BL v. PT3 p<0.0001)* suggesting strong memory recall and potential generalization of the fear response to the novel FR context.

**Figure 7.**
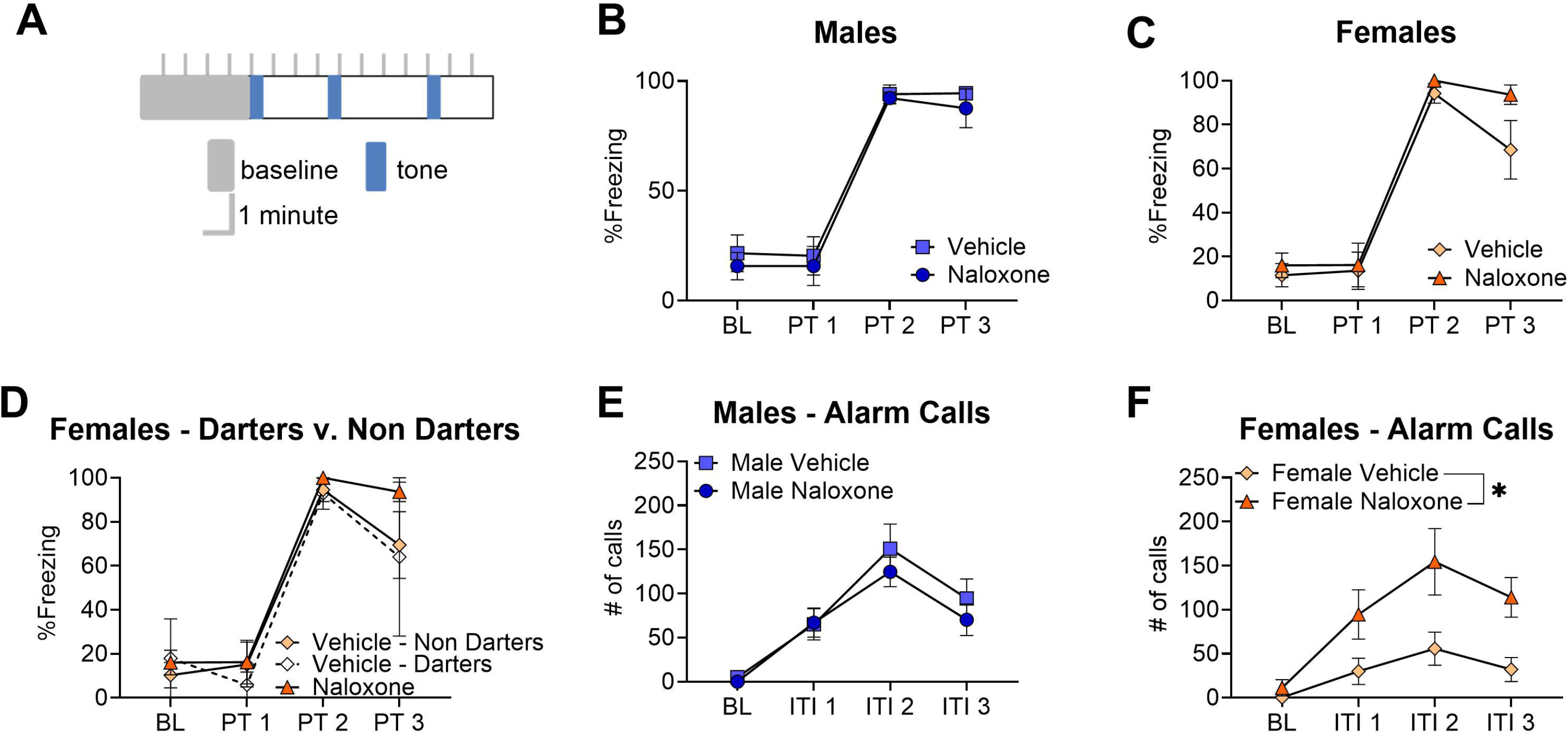
Naloxone increases ITI alarm calling during fear recall in females. **A**. Graphical representation of the timing of event related epochs during fear recall. **B-C**. Line graphs showing the percentage of time spent freezing during each pre-tone period (30s before tone presentation) for male and female animals injected with naloxone and injected with a vehicle prior to conditioning (Baseline: BL, PS: post-shock). **D**. Line graph showing the percent of time spent freezing during each pre-tone period by female groups with the vehicle injected group split into darters and non-darters. **E-F**. Line graphs showing the number of alarm calls emitted during each ITI period for male and female animals injected with naloxone and injected with a vehicle prior to conditioning (BL: baseline, ITI: intertrial interval). All graphs depict the mean ± SEM. Significant main effects are denoted with asterisks depicting degree of significance (1: p<0.05)

Males given naloxone did not differ from vehicle males in the number of alarm calls emitted during the ITI periods (Fig. 4E) (2-way RM ANOVA: no main effect of drug, F(1,18)=0.4579, p=0.51). Both male groups exhibited an increase in calling following the first tone presentation (*main effect of ITI period, F(2.170,39.06)=40.47, p<0.0001, Bonferroni post hoc comparison BL v. ITI1, ITI2, & ITI 3 all p<0.001*), suggesting strong fear memory recall.

Females given naloxone, however, did emit a higher number of alarm calls than vehicle females during the ITI periods (Fig. 4F) (*2-way RM ANOVA: main effect of drug, F(1,14)=8.532, p=0.011*). The number of emitted alarm calls from females given naloxone was similar to the number of alarm calls emitted by males. The results further cement that blocking MORs prior to conditioning alters patterns of USV emmittance in females during recall.

## [4] Discussion

The goal of this study was to examine the role of the endogenous opioid system in mediating several fear learning behaviors in both male and female rats. We found that administration of naloxone prior to fear conditioning altered behavioral responses in a sex- and behavior-dependent manner. While some effects were observed during conditioning, others emerged during recall, suggesting that MOR antagonism may differentially impact distinct components of fear learning and expression rather than producing a uniform change across behaviors.

We found that administration of naloxone prior to fear conditioning increased freezing in males. Previous work investigating the impact of naloxone on fear learning and memory has shown greater rates of freezing in males during testing, suggesting that it has a greater impact on memory consolidation or expression than learning (McNally et al., 2004; Yamasaki et al., 2024). Our results do not align with these findings, as we do not observe any difference in freezing rates during our recall test day. Differences in experimental design may partially account for these discrepancies. In the present study, naloxone was administered intraperitoneally at a dose of 5 mg/kg 15 minutes prior to conditioning, whereas McNally et al. (2004) used a 3 mg/kg subcutaneous injection and Yamasaki et al. (2024) used a 4 mg/kg dose administered 25 minutes prior to behavior. Variations in dose, route of administration, and timing relative to conditioning may alter the pharmacokinetic profile of naloxone and consequently influence its effects on fear learning and expression. However, because the increase in freezing that we observe during conditioning is concentrated to the baseline period and early tone presentations (tones 1 & 2), it suggests that MOR antagonism within our experimental perimeters may be having a greater impact on unconditioned responding rather than learning, as males given naloxone freeze at the same rate as vehicle males during middle and later tone presentations.

Naloxone affected unconditioned responding in females as well, leading to a decreased shock response velocity (the rate at which the animal moves directly following the footshock). Vehicle females had a significantly higher shock response velocity than both male groups, which aligns with our previous findings (Gruene et al., 2015; Colom-Lapetina et al., 2019; Mitchell et al., 2022). Typically, we observe velocities in females that are approximately 15 cm/s higher than their male counterparts (Mitchell et al., 2022), making the shock response rates of females given naloxone more similar to those typically observed in males. The female vehicle group was the only group to have any Darters during conditioning. Although Darters tend to have an increased shock response velocity relative to Non-darters (Gruene et al., 2015; Mitchell et al., 2022; Mitchell et al., 2024), the difference we observe between the female groups is not driven by darting alone, as the highest shock responses are not solely made up of the two darting animals. Impact of naloxone administration on activity during the 5-10 seconds after the shock has been previously observed (Fanselow, 1984), however they found increased mobility in response to the shock rather than a decrease.

Overall, this study had a lower number of Darters than we typically observe. Our average darting rates across studies is approximately 40% for females and 10% for males (Gruene et al., 2015; Mitchell et al., 2022; Mitchell et al., 2024). The highest rate of darting observed in any single group during this experiment was 17% for vehicle females during conditioning. This is likely due to random variation within this cohort of animals. However, these low numbers do not allow us to make meaningful comparisons between groups to determine the impact of naloxone on this behavioral response. Notably, the naloxone-treated female group in this study is the first female cohort we have observed with a complete absence of darters (Gruene et al., 2015; Colom-Lapetina et al., 2019; Laine et al., 2022; Mitchell et al., 2022; Huckleberry et al., 2023; Mitchell et al., 2024; Vincelette et al., 2026) making this finding particularly striking despite the overall low darting rates. Follow-up experiments will be required to fully determine whether MOR activity is necessary for the emergence of darting.

We observed no differences in ultrasonic vocalizations during fear conditioning. We have previously found equal numbers of shock response calls in males and females (Laine et al., 2022); however, we and others have typically observed a higher number of alarm calls in males (Graham et al., 2009; Schwarting et al., 2018; Laine et al., 2022), which we do not replicate here. In fact, we observed high rates of alarm callers in both female groups during conditioning.

During recall, however, a different pattern emerges. Females given naloxone had a much lower proportion of alarm callers, more in line with what is typically observed in females. And yet, those that did emit alarm calls produced a greater number of total calls than those in the female vehicle group across the entire testing session. They additionally emitted a greater number of calls during tone presentations compared to vehicle females, despite the greater amount of baseline calling observed in the vehicle group.

This difference in alarm calling during tone presentations may suggest that females given naloxone had enhanced consolidation relative to vehicle females, as previous findings have shown that naloxone can increase fear memory (McNally et al., 2004; Yamasaki et al., 2024). The difference in alarm calling between females given naloxone and those given a vehicle could also be related to the impact of naloxone on fear-conditioned analgesia (Fanselow, 1986; Fanselow & Helmstetter, 1988; Helmstetter, 1993). Blocking the activity of MORs prior to conditioning should attenuate the analgesic effect of the tone during later testing (Fanselow & Bolles, 1979b), thereby altering how the animals encode the shock—and, in turn, the cue. However, previous work suggests that females show little to no fear conditioned analgesia following conditioning (Llorente-Berzal et al., 2022; Stock et al., 2001) and those demonstrating increased consolidation have primarily measured memory through freezing (McNally et al., 2004; Yamasaki et al., 2024), where we do not observe differences between our female groups.

These findings raise the possibility that alarm calls and freezing reflect distinct components of fear learning and expression. Previous work (Zheng et al. 2025) has suggested that these behaviors may map onto separable processes, consistent with LeDoux and Pine’s “two-system framework” model of fear (LeDoux, 2022; LeDoux & Hofmann, 2018; LeDoux & Pine, 2016). Within this framework, freezing may represent a defensive response to potential threat, whereas alarm calls may reflect the subjective or affective experience of fear. It is therefore possible that naloxone administration prior to conditioning alters the subjective fear experience in females, without producing comparable changes in defensive responding. These findings also underscore the risk of interpreting freezing as a singular proxy for fear, as changes in affective state may occur independently from overt defensive responses.

It is worth noting, however, that this effect is only observed in approximately half of the females given naloxone, as the other half did not emit a single alarm call during the recall test despite the majority emitting alarm calls during conditioning. This intragroup variability raises important questions about what differences exist between these subpopulations of females. Our prior work investigating darting has already demonstrated the importance of examining individual variability within sex, highlighting that distinct behavioral strategies can emerge within female populations (see Gruene et al., 2015). It is possible that these findings represent an additional subdivision in how females engage in fear learning and memory consolidation.

Previous work has found that MOR expression can vary in the hippocampus, hypothalamus, and thalamus across stages of the estrous cycle (Piva et al., 1995; Kelly et al., 2003), which we did not examine here, and could explain the variability we observe.

Previous work has shown that naloxone administration can increase consolidation of fear memory for both cue and context (McNally et al., 2004), as well as increase non-cue freezing during conditioning, such as post-shock freezing (Fanselow & Bolles, 1979a). In the present study, however, this effect appears to be sex-specific. Females administered naloxone displayed elevated freezing during conditioning relative to naloxone-treated males, whereas males did not show comparable changes in freezing behavior. Interestingly, the work by Fanselow & Bolles (1979a) was conducted exclusively in females, raising the possibility that naloxone-induced increases in post-shock freezing may represent a uniquely female response pattern. Fanselow & Bolles observed dose-dependent increases in post-shock freezing across several naloxone doses (.5, 3, and 8 mg/kg), with the 5 mg/kg dose used in the present study falling within this effective range. This effect may reflect sex differences in the processing or long-term encoding of the aversive properties of the shock itself. Previous work has demonstrated that females are generally more sensitive to noxious stimuli (Wiesenfeld-Hallin, 2005), suggesting that MOR antagonism may have a greater impact on the subjective or physiological experience of the footshock in females than in males. Increased freezing during conditioning may also help explain the reduced shock-response velocity observed in naloxone-treated females.

Importantly, these alterations in shock responding also appear to influence subsequent fear-related behaviors during recall. In addition to increased alarm calling during cue presentations, naloxone-treated females also exhibited elevated alarm calling during the ITI periods. Notably, rates of alarm calling in naloxone-treated females were comparable to those observed in the male groups. More broadly, many of the behavioral changes observed in naloxone-treated females resulted in a more “male-like” behavioral phenotype. Together, these findings suggest that MOR signaling may be utilized differently in females during fear conditioning and recall, contributing to the sex-dependent patterns of defensive responding observed both here and in previous studies. It is also important to acknowledge that naloxone is not selective for mu-opioid receptors. Thus, while the present findings are discussed largely within the context of MOR signaling, contributions from delta- and kappa-opioid receptor systems cannot be ruled out. Future studies using receptor-selective antagonists or cell-type specific manipulations will be necessary to determine the extent to which these effects are specifically mediated by MOR signaling.

Overall, these findings suggest that MOR antagonism does not produce a uniform enhancement of fear learning, but instead differentially alters specific components of fear behavior in a sex-dependent manner. Across both males and females, naloxone appears to primarily influence unconditioned responding during conditioning; however, its effects in females during recall—particularly on alarm calling—point to a more selective role in shaping how fear is encoded and experienced. The dissociation we observe between freezing and alarm calling further supports the idea that these behaviors reflect distinct aspects of fear (Zheng et al. 2025), and that MOR signaling may preferentially modulate the affective or “subjective” component of the experience rather than defensive responding. Finally, the substantial variability observed within females highlights the importance of considering individual differences when interpreting sex effects and suggests that multiple behavioral strategies may exist within female populations. Together, these findings provide new insight into how the endogenous opioid system contributes to fear learning and expression and underscore the need to consider both sex and behavioral phenotype when investigating its role.

## Author Contributions

JF, MLL, & RMS conceptualized the study and designed the experiments. JF & MLL carried out the experiment and scored behavioral data. EMG conducted all statistical analysis and prepared figures. EMG & RMS wrote the manuscript.

## Data Availability

The data that support the findings of this study are available from the corresponding author upon reasonable request.

## Funding

Work funded by NIMH grant R01MH123803 to RM Shansky.

## Acknowledgements

We acknowledge Isabella Ravaglia, Kirti Gowda, Phoebe Shore, Gabrielle Kim-Levesque, Ece Ulgenturk, Emmett Bergeron, Vivika Sheppard, and Uma Nagella for technical support and assistance. Panel A in Figures 1-7 were created using Biorender.

## Notes

### Competing Interest Statement

The authors have declared no competing interest.

